# The genome of the biting midge *Culicoides sonorensis* and gene expression analyses of vector competence for Bluetongue virus

**DOI:** 10.1101/249482

**Authors:** Ramiro Morales-Hojas, Malcolm Hinsley, Irina M. Armean, Rhiannon Silk, Lara E. Harrup, Asier Gonzalez-Uriarte, Eva Veronesi, Lahcen Campbell, Dana Nayduch, Christopher Saski, Walter J. Tabachnick, Paul Kersey, Simon Carpenter, Mark Fife

**Affiliations:** The Pirbright Institute, Ash Road, Woking, Surrey GU24 0NF, UK; Rothamsted Insect Survey, Rothamsted Research, West Common, Harpenden, Hertfordshire, AL5 2JQ, UK; EMBL-European Bioinformatics Institute, Wellcome Trust Genome Campus, Hinxton, Cambridge CB10 1SD, UK; Bioinformatics group, Rothamsted Research, West Common, Harpenden, Hertfordshire, AL5 2JQ, UK; National Centre for Vector Entomology, Institute of Parasitology, Vetsuisse Faculty, University of Zürich, Zürich, Switzerland; USDA-ARS, Center for Grain and Animal Health Research, Arthropod Borne Animal Diseases Research Unit, 1515 College Avenue, Manhattan, KS 66502, USA; Clemson University Genomics Institute, Department of Genetics and Biochemistry, BRC #310, 105 Collins Street, Clemson, SC 29634, USA; Florida Medical Entomology Laboratory, Department of Entomology and Nematology, University of Florida, IFAS, 200 9^th^ St., SE, Vero Beach, FL 32962 USA

**Keywords:** Culicoides, biting midges, bluetongue virus, genome, genomics, transcriptomics, vector competence

## Abstract

**Background:** The use of the new genomic technologies has led to major advances in control of several arboviruses of medical importance such as Dengue. However, the development of tools and resources available for vectors of non-zoonotic arboviruses remains neglected. Biting midges of the genus *Culicoides* transmit some of the most important arboviruses of wildlife and livestock worldwide, with a global impact on economic productivity, health and welfare. The absence of a suitable reference genome has hindered genomic analyses to date in this important genus of vectors. In the present study, the genome of *Culicoides sonorensis*, a vector of bluetongue virus (BTV) in the USA, has been sequenced to provide the first reference genome for these vectors. In this study, we also report the use of the reference genome to perform initial transcriptomic analyses of vector competence for BTV.

**Results:** Our analyses reveal that the genome is 197.4 Mb, assembled in 7,974 scaffolds. Its annotation using the transcriptomic data generated in this study and in a previous study has identified 15,629 genes. Gene expression analyses of *C. sonorensis* females infected with BTV performed in this study revealed 165 genes that were differentially expressed between vector competent and refractory females. Two candidate genes, *glutathione S-transferase* (*gst*) and the antiviral helicase *ski2*, previously recognized as involved in vector competence for BTV in *C. sonorensis* (*gst*) and repressing dsRNA virus propagation (*ski2*), were confirmed in this study.

**Conclusions:** The reference genome of *C. sonorensis* has enabled preliminary analyses of the gene expression profiles of vector competent and refractory individuals. The genome and transcriptomes generated in this study provide suitable tools for future research on arbovirus transmission. These provide a significant resource for these vector lineage, which diverged from other major Dipteran vector families over 200 million years ago. The genome will be a valuable source of comparative data for other important Dipteran vector families including mosquitoes (Culicidae) and sandflies (Psychodidae), and yield potential targets for transgenic modification in vector control and functional studies.

## Background

Arboviruses (arthropod-borne viruses) are a taxonomically diverse group that include some of the most important emerging and re-emerging pathogens of wildlife, livestock and human beings worldwide [1–3]. Among arbovirus vectors, the majority of recent genomic studies have been carried out on the mosquitoes *Aedes* (*Stegomyia*) *aegypti* (L.) and *Aedes* (*Stegomyia*) *albopictus* (Skuse) due to their involvement in human to human transmission of a wide-range of arboviruses, their relative ease of colonization and recent technical advances in genome sequencing and annotation technologies [4], While these studies have led to major advances in control of the arboviruses these species transmit and our understanding of what drives susceptibility to infection [5–7], tools and resources for use with many other vector groups remain neglected.

To date, no genomic analyses have been carried out for vectors of non-zoonotic arboviruses that are pathogenic to livestock and wildlife. Among the most important of these vector groups are *Culicoides* biting midges (Diptera: Ceratopogonidae), species of which transmit internationally important arboviruses of ruminants, equines and deer [8–10]. In recent years, unprecedented outbreaks of *Culicoides*-borne arboviruses such as bluetongue virus (BTV) have inflicted huge economic losses on the livestock sector of Europe (e.g. estimate of US$ 1.4 billion in France in 2007) through clinical disease and accompanying restrictions against the movement of livestock imposed to limit virus spread [11,12], Globally, BTV outbreaks initiate non-tariff trade barriers restricting the movement of livestock and livestock germplasm, and cause a decrease in ruminant productivity in regions as diverse as India and the USA [13]. Globally, the economic impact of bluetongue has been estimated to US$ 3 billion [12].

*Culicoides* are notoriously difficult to culture under laboratory conditions and their small size of approximately 1.5mm body length renders them far from ideal subjects for transcriptomic and genome manipulation studies [14]. Only one of 14 confirmed vector species of *Culicoides* is currently colonized, *Culicoides sonorensis* Wirth and Jones [8,15]. *Culicoides sonorensis* colonies have already provided significant insights into the genetic basis of vector competence for BTV [16], a *de novo* transcriptome [17], and have been used to construct a physical map of the *C. sonorensis* genome which consists of four chromosomes [18,19].

Susceptibility to infection and transmission of arboviruses by *Culicoides* is determined in part by the heritability of barriers to virus dissemination following ingestion of the bloodmeal [13, 20, 21], Experimental evidence of a midgut infection barrier, a midgut escape barrier and a haemocoel dissemination barrier in *C. sonorensis* (formerly C. *variipennis* or *C. v. sonorensis* [22]) have been defined [23]. There is no evidence to date of salivary gland barriers preventing transmission in any species of *Culicoides* [24], as has been inferred in several species of mosquitoes [20]. Previous studies of the genetic basis for vector competence of *C. sonorensis* for BTV, using maternal inheritance properties, identified a 90 kd protein and used antibodies to this protein to isolate and characterize a cDNA clone encoding a glutathione S-transferase class delta enzyme [25]. Differential gene regulation in response to BTV infection, however, has not been investigated to date.

In this study, we sequence, *de novo* assemble, annotate and explore the first full genome of *C. sonorensis*. We then use transcriptomic analyses both to improve gene prediction within the genome build, and to elucidate differential gene expression associated with *Culicoides* competence for a BTV serotype 1 strain. The full genome sequence and transcriptome analyses are important resources for further studies of this phylogenetic group which is separated from the other major Dipteran vector families by at least 220 million years [26]. In addition to being a valuable resource for comparative study of vector phylogenomics, the provision of *Culicoides* genomes is of interest as a group that demonstrates several unique features, including a significant capacity for long-distance dispersal by semi-passive flight and the ability to reach huge population density under suitable conditions. In addition, their close association with livestock raises questions concerning both host preference and vector competence for arboviruses. Many of these questions can be readily addressed by understanding genetic diversity within populations which will be enhanced by the provision of comparative genomic data, but also in the long term by yielding targets for transgenic modification. The present genome provides a resource to facilitate all these studies.

## Methods

### Samples

All *C. sonorensis* used in this study originated from the ‘AA’ colony which was originally established in 1955 at the Kerrville, Texas laboratory of the US Department of Agriculture [27], Since 1969, this colony has been maintained at The Pirbright Institute without any additional outbreeding [28]. Within this period, two selection bottlenecks of ≤10 individuals were performed to increase susceptibility to BTV serotype 4 and African horse sickness virus (AHSV) serotype 9 infection (Mellor, Pers Comm). Following the second selection in the 1990’s [29], the colony strain was renamed as PIR-s-3. No attempt was made to further reduce heterozygosity prior to using individuals from the PIR-s-3 line in this study. All *C. sonorensis* used were 3-4 days old and had not been fed sucrose following emergence from pupae.

### DNA extraction

The genomes of *C. sonorensis* males and females were sequenced in separate. Adults were separated by sex under a stereomicroscope and stored in 95% ethanol at room temperature prior to use. In order to obtain sufficient amount of DNA for genomic sequencing, DNA was extracted from 375 males in three pools of 100 and one pool of 75 individuals, and from 150 females in one pool of 100 and one pool of 50 individuals. Pooled samples were homogenised twice for 30 seconds in 75 μl of phosphate buffered saline using a Tissuelyser™ (Qiagen, UK) at a frequency of 25 Hz/s and a 3mm stainless-steel ball bearing (Dejay Distribution Ltd, UK) within 2ml screw-topped tubes. Genomic DNA extraction was conducted using gravity-flow anion-exchange tips (Qiagen’s Genomics Tip 20/G) with extraction of DNA up to 150kb in size and a maximum of 20μg of product A user-developed protocol for mosquitoes and other insects was followed (see Supplementary Methods). The DNA obtained from each extraction was pooled by sex and the concentration of the resulting samples was evaluated using a Qubit^®^ 2.0 Fluorometer (ThermoFisher Scientific, UK) with the Qubit^®^ DNA BR Assay Kit (ThermoFisher Scientific, UK). The integrity of the genomic DNA was visualized using a 5 μl sample on a 1% (*w/v*) agarose gel.

### Genome sequencing and *de novo* assembly

8 – 9 μg of high molecular weight (HMW) genomic DNA were used to sequence the genome of male and female *C. sonorensis* separately. Two paired-end (PE) libraries of insert sizes 200 bp and 400 bp and a mate-pair (MP) library with an average fragment size of 4.4 kb were sequenced with 150 bp (200 bp and 4.4 kb libraries) and 100 bp (400 bp library) paired-end from both female and male pools were run on Illumina HiSeq 2000 and 2500 sequencing systems. Library construction and sequencing was performed at The Earlham Institute (Norwich, UK). Read pair quality was assessed with FastQC vO.lO.l. Results suggested that 6 bp should be removed from the start of each read, and in the case of MP data the last 40-50 bp had unexpectedly high levels of k-mer duplication. Both PE and MP reads were filtered to limit their size to lOObp after removal of the first 6 bp. MP data was processed with NextClip vl.O [30] to select reads which contained both ends of the original fragment. This created a dataset of reads with variable length, which were subsequently limited to 100bp.

PE and MP libraries were initially assembled using Velvet v1.2.08 [31] with a k-mer length of 91. The N50 of the assembly was maximized using an expected coverage of 27, a coverage cut-off of six, and with scaffolding option set to ‘no’. This initial assembly was filtered to remove short (<4kb) and repeated contigs (> 90% sequence ID to a larger contig) using BLAT v34 and a custom script to count identical bases, considering any overlaps between matches. The assembly was then scaffolded using SSPACE v2.0 [32] and the 400bp PE library, using three passes of decreasing stringency. Any remaining contigs smaller than 500bp were removed (9913 total). The resulting assembly was then scaffolded a second time using the 4.4kb MP library with SSPACE v2.0 (Figure 1). This assembly was assessed using FRCbam vl.O [33] and Reapr vl.O.15 [34], and the output from Reapr was processed with GapFiller vl.ll [35]. The final assembly was assessed using FRCbam vl.O to confirm that an improvement in the reported error rate had occurred. Genome size was estimated using the number of total trimmed nucleotides which were used in the assembly divided by the maximum of the per-position coverage frequency distribution, after reads were mapped back to the assembly using BWA-MEM [36]. Kmer spectra were assessed using KAT (https://katreadthedocs.io/en/latest). Redundant contigs were identified using Redundans [37].

**Fig. 1.**
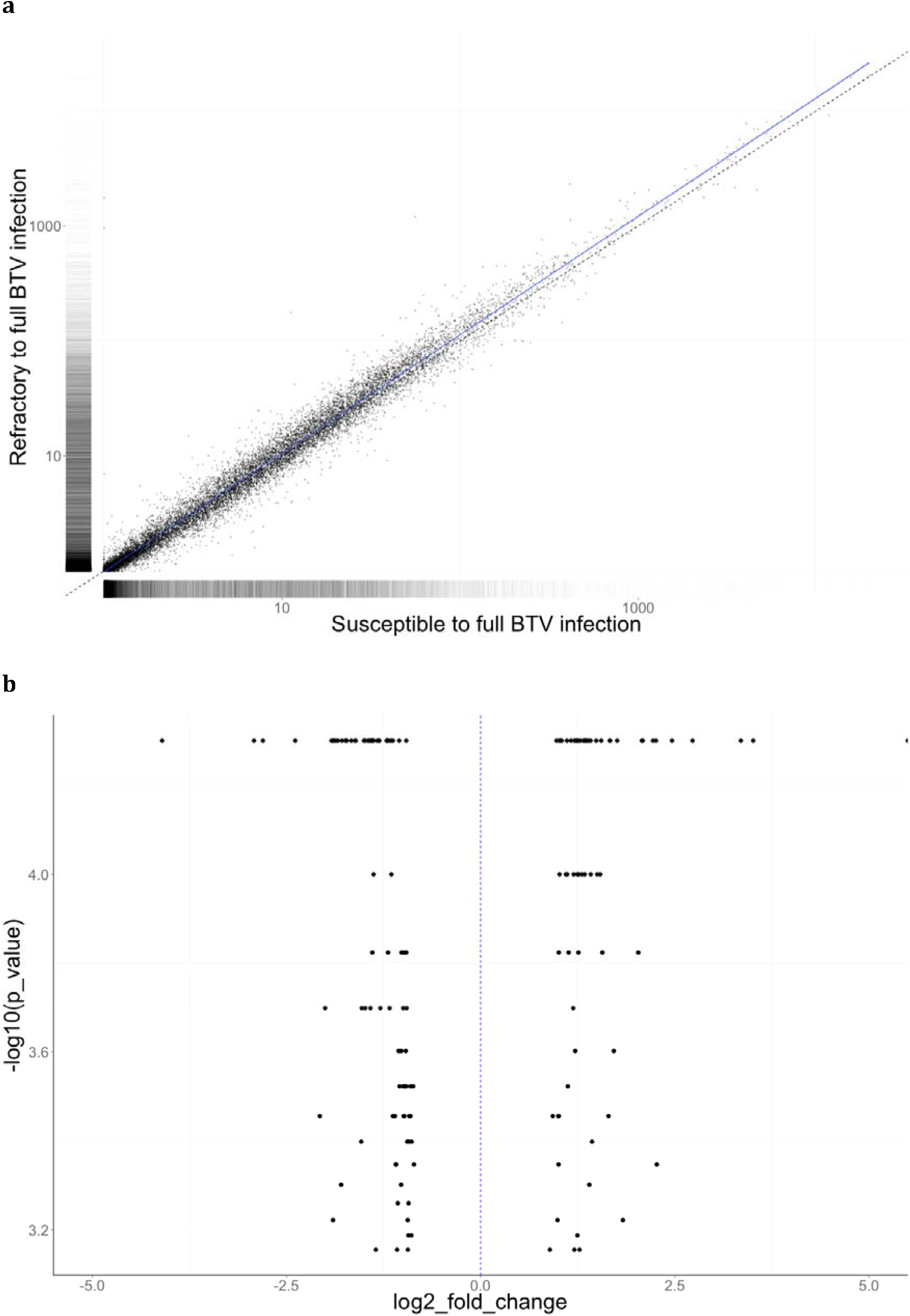
Differential gene expression analyses between females susceptible to full BTV infection and refractory females, **a** Scatterplot sof the pairwise comparison of the gene expression levels between the two phenotypes. The average gene expression across all genes is displayed by the blue line, **b** Volcano plot displaying fold changes in expression of the 165 differentially expressed genes between *Culicoides sonorensis* that are susceptible to infection and those that are refractory to full infection with BTV.

Data (reads, annotation and assembly] have been deposited in the ENA database under the accession number PRJEB19938. The genome used in this present study is the latest, v1.

### Gene model prediction and annotation

Prediction and annotation of the genome assembly was conducted using MAKER v2.31.6 [38]. The genome annotation was done using the transcriptome data reported in this study and transcriptome data from previous studies on *C. sonorensis* [17, 39]. In addition, we used the genome annotations of ten species of *Anopheles, Ae. aegypti, Belgica antarctica* Jacobs, *Culex quinquefasciatus* Say *Drosophila melanogaster* Mg., *Lutzomyia longipalpis* (Lutz & Neiva, 1912] and *Phlebotomus papatasi* (Scopoli] (see Supplementary Methods for the species and genome assembly versions used]. SNAP v2006-7-28 [40] and AUGUSTUS v2.5.5 [41] were used to predictαł) *initio* gene models. SNAP v2006-7-28 was trained using 500 models and AUGUSTUS v2.5.5 using the best 1000 models extracted from the initial MAKER v2.31.6 output, as recommended in the software documentation. Coverage of conserved proteins was assessed using the CEGMA v2.4 [42] and BUSCO v1 [43] pipelines. Analysis of Gene Ontology annotation was performed with Blast2GO v3 [44], The orthology of the annotated genes was established using the Ensembl Compara Gene Trees pipeline [45], using the genomes of *C. sonorensis* and 17 other arthropod species as input (see Supplementary Metods]. Genes were functionally annotated using InterProScan, which computationally identifies the presence of domain, motif or family signatures within a protein sequence and infers functional descriptors (taken from the Gene Ontology (GO)).

### Infection experiments with bluetongue virus

The virus used to infect *C. sonorensis* was a western topotype strain of BTV serotype 1 (GIB2007/01) [46]. Virus stock was mixed 1:1 with defibrinated horse blood (TCS Biosciences, Buckingham, UK) with 6.2 Logio TCIDso/ml being the final infectious dose. All blood-feeding of *C. sonorensis* was conducted using a Hemotek membrane-based system (Discovery Workshops, Accrington, UK). A total of 150 and 145 *C. sonorensis* in two replicates were fed on the blood:BTV-1 suspension and survived the extrinsic incubation period of eight days at 25 °C. During this period of incubation, 10% (w/v) sucrose solution was offered via a cotton wool pad. Each individual was then decapitated using disposable needles and the head and remainder of the body of each were stored separately in RNAlater™ (Thermofisher Scientific, UK) at −20 °C.

### RNA extraction

RNA extraction from *C. sonorensis* was done following the TRIzol^®^ (ThermoFisher Scientific, UK) protocol (see Supplementary Methods). In brief, samples were homogenised in 100 μl of Schneider’s *Drosophila* media (Gibco™, Thermofisher Scientific, UK) and RNA was extracted using TRIzol^®^ and chloroform, followed by a precipitation with Isopropanol. The integrity of the RNA was analysed using a Bioanalyser 2100 and the concentration was estimated on a Qubit^®^ 2.0 Fluorometer (ThermoFisher Scientific, UK) with the Qubit^®^ DNA BR Assay Kit (ThermoFisher Scientific, UK).

RNA was extracted from individual heads of *C. sonorensis* fed with blood:BTV-1 as described for the bodies (Supplementary Methods). The corresponding bodies (abdomen and thorax) of the BTV-1-positive and BTV-1-negative heads were pooled separately for RNA extraction. In addition, the RNA from 100 *C. sonorensis* fed on horse blood three days post-emergence and left without access to sucrose for eight days, and from 100 *C. sonorensis* fed on a 10% (w/v) sucrose solution from 3-8 days following emergence, were also extracted and their transcriptomes sequenced as controls.

### Quantification of infection with BTV-1

We used RT-qPCR to detect and quantify the presence of disseminated BTV-1 viral RNA in the heads of blood:BTV-1 fed *C. sonorensis*. For this, the RNA was first reverse-transcribed into DNA using the ProtoScript II First Strand cDNA Synthesis kit (New England Biolabs) (see Supplementary Methods for a detailed protocol). A reaction with no enzyme was included as a no-RT negative control. The resulting cDNA was used as template in RT-qPCRs for the detection of BTV using SYBR Green assays (Supplementary Methods). The reactions were performed using the BTV specific primers BTVuni 291-311F 5’ GCTTTTGAGGTGTACGTGAAC 3’ and BTVuni 381-357R 5’ TCTCCCTTGAAACTCTATAATTACG 3’ [47], No-template and no-RT reactions were included as negative controls. To verify that negative reactions were not due to a lack of cDNA in the sample, we also performed RT-qPCR reactions for each sample using *C. sonorensis* specific primers for the Vacuolar ATPase gene (Vac-ATPase forward 5’ GCTGCTGCTGCCATCATTTT 3’ and Vac-ATPase reverse 5’ CCGGTCGCATCACTGACATA 3’). All reactions were performed in duplicate. Identification of samples with and without BTV-1 RNA was done using the quantification cycle (C_q_) values of the reactions. For the purposes of this study, we considered *vector competent* individuals to be those with disseminated infections that include replication in the head capsule, as this has been demonstrated to allow the isolation of BTV-1 [46]. We refer as *refractory individuals* to those without detectable BTV after a suitable incubation period (eight days), those with BTV midgut infections that had not disseminated within the insect, those where the process of dissemination had not been completed (i.e. those with haemocel dissemination barriers, midgut escape and infection barriers) and those still retaining inactivated BTV following the infective bloodmeal [24], In the case of this study, we used a C_q_ value of 27 to differentiate between vector competent and vector refractory (see Results).

### Transcriptome sequencing and analyses

In total, eight transcriptomes were sequenced: two biological replicates each for i) BTV-competent; ii) BTV-refractory; iii) blood-fed; and, iv) sucrose-fed. Library construction and sequencing was performed at Edinburgh Genomics (The University of Edinburgh, Scotland). A total of 2.5 μg of RNA from each sample was used to construct TrueSeq libraries, which were sequenced using 50 base paired-end (PE) reads on an Illumina HiSeq 2500. Each library was run in two lanes.

The quality of the raw sequence data was analyzed in FastQC vθ.11.2. Reads were aligned to the genome assembly (Illumina assembly, without removal of the redundant contigs) with TopHat v2.0.6 [48] using the gene model annotations generated from the genome as reference. The resulting bam files were assembled with Cufflinks v2.2.1 [49] and merged into a single transcriptome with Cuffmerge v2.2.1. Quantification of gene and transcript expression and comparison of the expression levels was performed with Cuffquant v2.2.1 and Cuffdiffv2.2.1, respectively. The Cuffquant results for each transcriptome (two biological + two sequencing lane runs) were grouped per experimental conditions (blood-fed, sucrose-fed, vector competent and vector refractory) to compare the expression profiles between these different experimental conditions with Cuffdiff. Differential expression analyses were then explored with the R package CummeRbund v2.9.3 [50]. Differentially expressed (DE) genes with a significant change in expression level were identified using an *α* value of 0.05, which establishes the filtering value of the multiple-testing corrected q-values. Identification and functional classification of the DE genes was done using BLAST and InterProScan searches in Blast2Go version 4.0.2. Enrichment and gene set enrichment analyses were performed on the sets of DE genes between the different conditions using the interface provided in Blast2Go to the Fisher’s Exact Test implemented in FatiGO [51] and the Gene Set Enrichment Analysis (GSEA) package [52], To perform the GSEA, all genes were ranked per the logarithmic fold-change, removing the infinite values. Transcriptome data have been deposited in the ENA database under the accession numbers ERR2171964-ERR2171979.

### Validation of differentially expressed genes

Quantitative reverse transcription PCR (RT-qPCR) was used to validate the change in expression of four differentially expressed (DE) genes between the vector competent and refractory pools of *C. sonorensis*. These DE genes were selected based on previous identification in vector competence studies of *C. sonorensis* or because they have a direct functional link with antiviral response. The genes were: a gene sharing 73% identity with the antiviral helicase *ski2* of *Cx. quinquefasciatus (ski2;* XP_001845019); a gene sharing 63% identity with *glutathione S-transferase* (*gst1* XP_001654620), a gene sharing 96% identity with *glutathione S-transferase-1* from *C. sonorensis (gst-1?* AAB94639) and a gene sharing 49% identity with a gene encoding a *Toll* protein in *Acyrthosiphon pisum* (Hempitera: Aphididae) (XP_001948700). The RNA used to validate the change in expression were the two biological replicas of vector competent and refractory *C. sonorensis* transcriptomes sequenced.

Expression levels were normalized against three reference genes whose stability was ranked using three approaches, BestKeeper [53], geNorm [54], and NormFinder [55] (Supplementary Table S3). Gene-specific primers and probes were designed using Primer3Plus [56] (Supplementary Table S4; Supplementary Methods). Primers were compared against the *C. sonorensis* genome using BLAST to ensure specificity to the target region. Amplification efficiency was tested by RT-qPCR with hydrolysis probes by generating a standard curve using triplicate technical replicates of five-fold serial dilutions of RNA template from teneral female *C. sonorensis* specimens. All RNA templates were treated with Turbo DNA-Free™ Kit (ThermoFisher Scientific, UK) prior to RT-qPCR to remove any DNA from the sample following the manufacture recommended protocol. Starting template total RNA concentration was evaluated using a Qubit^®^ 3.0 Fluorometer (ThermoFisher Scientific, UK) with the Qubit^®^ RNA HS Assay Kit (ThermoFisher Scientific, UK).

cDNA synthesis and quantitative amplification was done in one reaction using the Superscript^®^ III One-Step RT-qPCR System with Platinum^®^ Taq DNA polymerase (ThermoFisher Scientific, UK) (see Supplementary Methods for a detailed protocol). Reactions were performed following the fast cycling programme as described in the manual (Supplementary Methods). PCR amplification efficiency of each primer-probe-target combinations was calculated via the linear regression of G, as the log2 of the relative RNA template concentration using the ggplot2 package [57] in R. Reactions which exhibited amplification efficiency between 90-110% with an R^2^ value of ≥0.98 were considered valid. Relative expression levels for the gene of interest between vector competent and refractory *C. sonorensis* were calculated using the ΔΔCq method [58].

### Molecular evolution analyses

Homologues of the *ski2* and *gst-1* genes in other insect species were identified using BLAST in Blast2Go. The most significant BLAST hits from *Ae. aegypti, Cx*.

*quinquefasciatus* or *An. gambiae* were then used to identify the orthologues in other vector species in VectorBase [59]. We verified that the sequences used started with Methionine and ended with a stop codon, whenever possible. The protein sequences identified were aligned using the mode expresso of T-Coffee [60]. Poorly aligned regions (positions with score 0-6) were trimmed from the alignment and the resulting alignments were used in the evolutionary analyses (see Supplementary Methods for more details and commands used to align the sequences; Supplementary Alignment files which include the sequences before and after alignment in fasta format). Genetic distances and the diversity were estimated using the software MEGA v7.0.26 [61]. Phylogenetic analyses were performed in MrBayes v3.2 [62] using the protein alignment as input and the models of evolution that best fitted our alignments were identified using ProtTest 3.4.2 [63]. Two independent analyses of two million generations with four chains (one cold and three heated) were conducted. Trees and parameters were sampled every 50^th^ generation and the first 10,000 were discarded. The remaining trees were used to estimate the consensus and Bayesian posterior probabilities of each branch. The s*ki2* from *Saccharomyces cerevisiae* Meyen ex E.C. Hansen and the *gst-1* from *Pediculus humanus* L. were used as outgroups.

## Results

### Genome sequencing, construction and annotation

Assembly of the Illumina reads resulted in a genome of 197,417,436 bp assembled in 7,974 scaffolds. In contrast, the total genome size estimated using the method of Schell *et al*. [36] (from the maximum depth of read coverage) was 294.45 Mb (Supplementary Table Sll). The N50 of the genome assembly was 89,502 and the proportion of Ns was of 2.63% (Table 1). Analysis with Repeat Modeller [64] identified a total of 793 repeat regions encompassing —14% of the genome. When low-complexity regions are included, the total repeat content is 29.7% including 2 Mb of Type II transposons and 5 Mb of Type I transposons (mostly LINE elements). The GC content of the *C. sonorensis* genome is 28% in the complete assembly and 34% in the exons (Table 1). This is low when compared to the closest fully sequenced relative to *C. sonorensis* (*Belgica antarctica* (Diptera: Chironomidae): total GC content = 39%) and a range of other Diptera genomes (Table 1).

**Table 1.** A comparison of genome characteristics of the *C. sonorensis* genome with other selected Diptera species

| **Parameter** | ***Culicoides***<br>***sonorensis*** | ***Aedes aegypti***<br>**L3** | ***Aedes aegypti***<br>**L5** | ***Anopheles***<br>***gambiae***<br>**AGAM P4** | ***Drosophila***<br>***melanogaster***<br>**BDGP6** | ***Belgica antarctica***<br>**GCA_000775305.1** |
| --- | --- | --- | --- | --- | --- | --- |
| **Genome size (Mb)** | 197.4 | 1,384.1 | 1,362.0 | 278.3 | 143.7 | 89.7 |
| **Number of contigs** | 15,810 | 36,205 | 4,901 | 16,825 | 2,442 | 22,492 |
| **Contig N50** | 30,774 | 82,500 | 10,861,303 | 85,555 | 19,488,218 | 13,551 |
| **Contig N90** | 5,567 | 15,283 | 74,389 | 5,600 | 666,663 | 1,865 |
| **Number of scaffolds** | 7,974 | 4,757 | 4,616 | 10,523 | 1,870 | 4,997 |
| **Scaffold N50** | 89,077 | 1,547,048 | 409,777,670 | 49,364,325 | 25,286,936 | 98,164 |
| **Scaffold N90** | 14,215 | 324,062 | 83,687 | 2,684 | 23,513,712 | 18,418 |
| **Ns (%)** | 2.8 | 5.3 | 0.0 | 7.6 | 0.8 | 0.9 |
| **Maximum contig length** | 552,397 | 685,587 | 71,953,859 | 808,130 | 27,905,041 | 131,402 |
| **Repeats + low complexity**<br>**regions** | 29.7% | 73% | 78% | 25% | 23% | 4% |
| **GC content** | 28.4% | 38.3% | 38.2% | 44.3% | 42.0% | 38.9% |
| **GC exons** | 34.3% | 46.8% | 46.2% | 54.6% | 49.1% | 47.1% |
| **Protein-coding genes** | 15,629 | 15,796 | 14,626 | 13,008 | 13,918 | 13,510 |
| **Mean protein-coding gene length** | 5,040 bp | 18,126 bp | 46,920 bp | 6,442 bp | 6,960 bp | 2,555 bp |
| **Mean exon length (protein coding genes)** | 480 bp | 460 bp | 494 bp | 438 bp | 538 bp | 325 bp |
| **Mean intron length (protein-coding genes)** | 632 bp | 23,265 bp | 10,456 bp | 1,008 bp | 1,147 bp | 213 bp |
| **Mean introns per transcript (in protein-coding genes)** | 6.7 | 5.1 | 7.9 | 5.5 | 6.9 | 5.3 |
| **Maximum splice isoforms/protein-coding gene** | 19 | 20 | 50 | 20 | 75 | n.a. |

Contigs refer to the contiguous sequences assembled without Ns; scaffolds refer to the final top level sequence assembly used for annotation (they contain Ns).

Analyses on the completeness of the assembly identified 98.79% (100% including partial matches) of 248 Core Eukaryotic Genes Mapping Approach (CEGMA) genes and over 97% of the Benchmarking Universal Single-Copy Orthologs (BUSCO) gene set for Insecta (97.1%), Arthropoda (97.7%) and Metazoa (97.3%), with additional genes from these sets (0.7-0.9%) found in fragmentary form. Of these genes, 65.5% (Metazoa)-66.3% (Insecta, Arthropoda) were found in single copy, compared with 89.3% of Insecta BUSCO genes that were present in single copy in th*eAedes aegypti* assembly, and a higher number for some other assemblies (Supplementary table S12). This indicates that there is some level of duplication within the *Culicoides* genome assembly, which could be due to a variation among/within the sequenced genomes and the separate representation within the assembly of alternative alleles.

Annotation of the Illumina assembled genome resulted in 15,629 protein-coding genes and 21,336 transcripts, which is comparable to the number of protein-coding genes in other species of Diptera (Table 1). The mean gene, intron and exon lengths in *C. sonorensis* are 5,040 bp, 828 bp, and 480 bp, respectively. The average number of introns per gene in *C. sonorensis* is 6.7. A total of 2,991 genes have been estimated to be alternatively spliced and the maximum number of splice variants per gene (19) is comparable to *Ae. aegypti* and *An. gambiae*.

To assess redundancy, we generated the K-mer spectra of the assembly and removed potentially redundant contigs using Redundans. This generated an alternative assembly of 156 Mb, with 3,839 contigs, and an N50 scaffold length of 109,184 bp. Coverage of the BUSCO data sets was not changed, with values for Insecta, Arthropoda, Metazoa and Eukaryota within +/-0.1% of the original assembly. The proportion of duplicate BUSCOs, however, was reduced from 30.8% (unreduced Illumina assembly) to 12.1%-13.2% (reduced assembly) (Supplementary Table S13), confirming the presence of BUSCO gene family members within the redundant contigs.

Homology to genes annotated in other species was detected for 93% of the genes annotated in *C. sonorensis*, with an average of 19 homologous genes and nine orthologues identified per gene; these numbers are comparable to the other species used in the analysis. Over 90% of the *C. sonorensis* genes were linked to their homologues via the ancestral nodes of Arthropoda (6,609 genes), Neoptera (6,361 genes) or Diptera (1,225 genes).

InterProScan was used to detect protein domains and assign GO terms to genes. Overall, a similar number of domains and high-level GO terms were associated with at least one gene as in most other sequenced insects (Supplementary table S14), although a larger number of specific GO terms tend to have been applied in genomes that have been manually annotated for gene function. In total, 46,329 GO term assignments were made. This compares to 45,600 assignments for An. *gambiae*, and 103,897 assignments for *D. melanogaster* (in which many terms have been manually assigned based on the literature, in addition to automatic assignments). Of 149 high-level categories in the GO (“GO Slim”), 137 were present in *C. sonorensis* as opposed to 140 categories represented in the genome of *An. gambiae* and 144 in *D. melanogaster*. GO terms assigned to more genes in *C. sonorensis* than either *An. gambiae* and *D. melanogaster* include: ion binding (2,869 genes, as opposed to 2543 in *An. gambiae* and 2,349 in *D. melanogaster*), oxidoreductase activity (703 genes, as opposed to 622 in *An. gambiae* and 632 in *D. melanogaster*), kinase activity (367 genes as opposed to 315 *in An. gambiae* and 363 in *D. melanogaster*), hydrolase activity acting on glycosyl bonds (112 genes as opposed to 105 in *An. gambiae* and 107 in *D. melanogaster*) and tRNA metabolic process (101 genes as opposed to 86 in *An. gambiae* and *D. melanogaster*). These results suggest an increase in the number of genes encoding these functions in *C. sonorensis* compared with other species, although the possibility that such genes are selectively over-represented in the current *C. sonorensis* assembly cannot be ruled out.

To identify tandemly arrayed genes, we looked for genes that contained common domain architectures (as defined by the set of InterPro-defined domains found within each gene) and were located within 100,000 nucleotides of each other. 1,281 genes were found in such arrays. The longest such array contained 10 copies of a gene containing a chitin-binding domain. The longest array of a protein with this architecture comprises just two copies in *D. melanogaster*, and there are no such arrays *\x\Ae. aegypti* or *An. gambiae*, although there are various longer arrays of proteins containing this domain alongside others in these species (Supplementary Table S15). The second longest such array contains 9 copies of a gene encoding a protein with arrestin and immunoglobulin domains; the homologous proteins are found in 10 tandem copies in *Ae. aegypti* and *D. melanogaster*, and four tandem copies in *An. gambiae* (Supplementary Table S15).

### Transcriptome sequencing and analysis

Paired-end sequencing of transcriptomes resulted in an average of 50 million reads with a length of 50 bp for each sample sequenced. Phred scores exceeded 34 in all cases. The transcriptomes from the different conditions were merged with Cuffmerge prior to a comparative analysis of their gene expression profiles. The number of genes on the merged transcriptome is 17,263 and the number of transcripts is 35,813. Of the genes present in the transcriptome, 15,630 had already been annotated in the genome. 1,816 transcripts, mapping to 1,635 loci, were novel. The proportion of gene loci with a single transcript is 62.7% (10,826), but 93.3% among the genes that had not been annotated on the genome. BLAST searches against NCBI’s Arthropod non-redundant (nr) protein database resulted in 92.4% of the isoforms (33,095) and 90% of the genes matching to a database entry (15,268 of the genes had one or more hits).

### Differential expression of genes between competent and refractory *Culicoides sonorensis*

Eighty-five (56%) *C. sonorensis* in replicate one and 53 (37%) in replicate two produced C_q_ values ≤ 27 (Supplementary Figure S1). We used the C_q_ value 27 as a cut off for differentiating between vector competent and refractory individuals as the distribution of values was bimodal and 27 was the local minimum (Supplementary Figure S1). Sixty-five (44%) *C. sonorensis* in replicate one and 92 (63%) in replicate two produced C_q_ values of > 27 or no C_q_ value. Replicate one had a significantly greater proportion of disseminated infections than replicate two (Fishers exact test; df=1,P = 0.0007). Three and nine individuals from the first and second experiment replicates were false negatives, that is, samples that were negative for BTV-1 but that also failed to amplify the *C. sonorensis* genes. This indicated that the RNA extraction failed and were discarded from further analyses. Transcriptome studies were therefore conducted using 147 and 136 female *C. sonorensis*, from the two biological replicates of the infection study, respectively.

Gene expression profiles showed that more genes had their expression upregulated in refractory *C. sonorensis* than in competent individuals (Fig. 1a). A total of 165 genes demonstrated significant differential expression (DE) (Fig. 1b; Supplementary Table S1) of which 123 (75%) had been annotated and 42 (25%) represented novel genes (Supplementary Table S2). A total of 94 (57%) were upregulated in refractory individuals, while 71 were downregulated (43%) (Supplementary Table S1). 120 of the 165 genes showing DE had one or more BLAST hits in the non-redundant (nr) protein database from NCBI (Supplementary Table S2) and 88% of these corresponded closely to genes from other members of the suborder Nematocera. Of the 165 genes, 63 were associated with gene ontology (GO) terms (Supplementary Figure S2). However, enrichment analysis showed no significant results.

*Ski2* (XLOC_006435) and *glutathione S-transferase-1* (CSONOH559) showed an increase in expression levels in the females that were resistant to full dissemination, the log2 fold change being −1.10 (P = 0.00035; q-value = 0.00381) and −1.07 (P = 0.00055, q-value = 0.00555), respectively (Fig. 2; Supplementary Table S1). In contrast, the *Toll-like* (XLOC_000522) gene and the gene matching *glutathione s transferase* (CSON010973) were expressed at higher levels in the samples that were susceptible to a fully disseminated BTV infection (log2 fold change 1.21, P = 0.00005 and q-value = 0.00072) and (log2 fold change 2.07, P= 0.00005 and q-value = 0.00072) (Fig. 2; Supplementary Table S1).

**Fig. 2.**
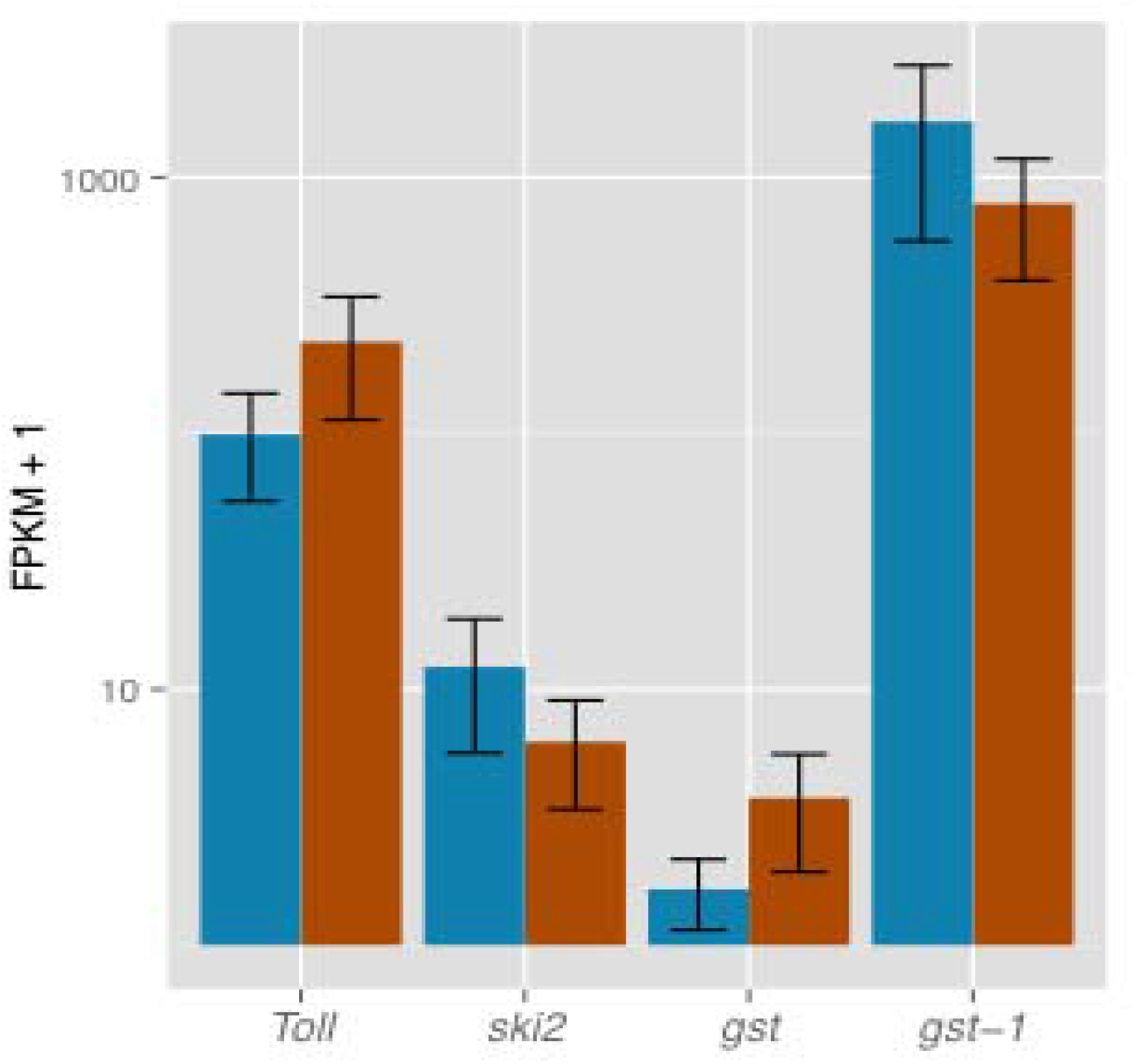
Expression bar plots for candidate genes identified as differentially expressed at significant level between vector competent (orange bars) and refractory (blue bars) samples to full BTV infection. *Toll* corresponds to the *Toll*-like gene, *ski2* is the antiviral helicase *ski2, gst* is *glutathione s transferase*, and *gst-1* is the *glutathione s transferase-1*.

The direction and degree of change in expression levels of the four genes identified as potentially influencing *C. sonorensis* competence for BTV (*Ski2, gst-1, gst* and To//-like) was confirmed using RT-qPCR and the same samples. Housekeeping genes used as reference were *CytB5, rpl13* and *rps8*, which showed the highest level of stability of those tested (Supplementary Table S3 and Table S4). The direction of up and down regulation of the genes and the log-2 fold change was similar in all cases as that detected in the RNAseq data (Supplementary Table S5).

### Molecular evolution of *ski2 antiviral helicase* and *glutathione s transferase-1*

The most significant BLAST hit for the *C. sonorensis ski2* was the gene from *Cx. ąuinąuefasciatus* (XP_001845019.1 = CPIJ003293). There are 37 genes identified as orthologous to *Cx. ąuinąuefasciatusski2* in 36 species in VectorBase. 35 are 1-to-1 orthologues and there are two orthologous genes *mAn. maculatus* (Supplementary Table S6a). The mean evolutionary diversity estimated using JTT model with gamma distribution value α = 0.5 (as estimated with ProtTest) and pairwise deletion across the sequences from Diptera species (no outgroups included) was 0.513, and pairwise distances between protein sequences ranged between 0.000 (*An. coluzzi* v. *An. gambiae*) and 1.198 (*G. fuscipes fuscipes* v. *An. maculatus*). The distance of the ski2 protein between *C. sonorensis* and mosquitoes was 0.752, with sandflies was 0.834, and with Brachycera had a mean distance of 0.928. The mean evolutionary distance within the Nematocera was 0.272, and 0.171 within the Brachycera. The evolutionary relationship of this protein is consistent with species-level phytogeny (Fig. 3a).

**Fig. 3.**
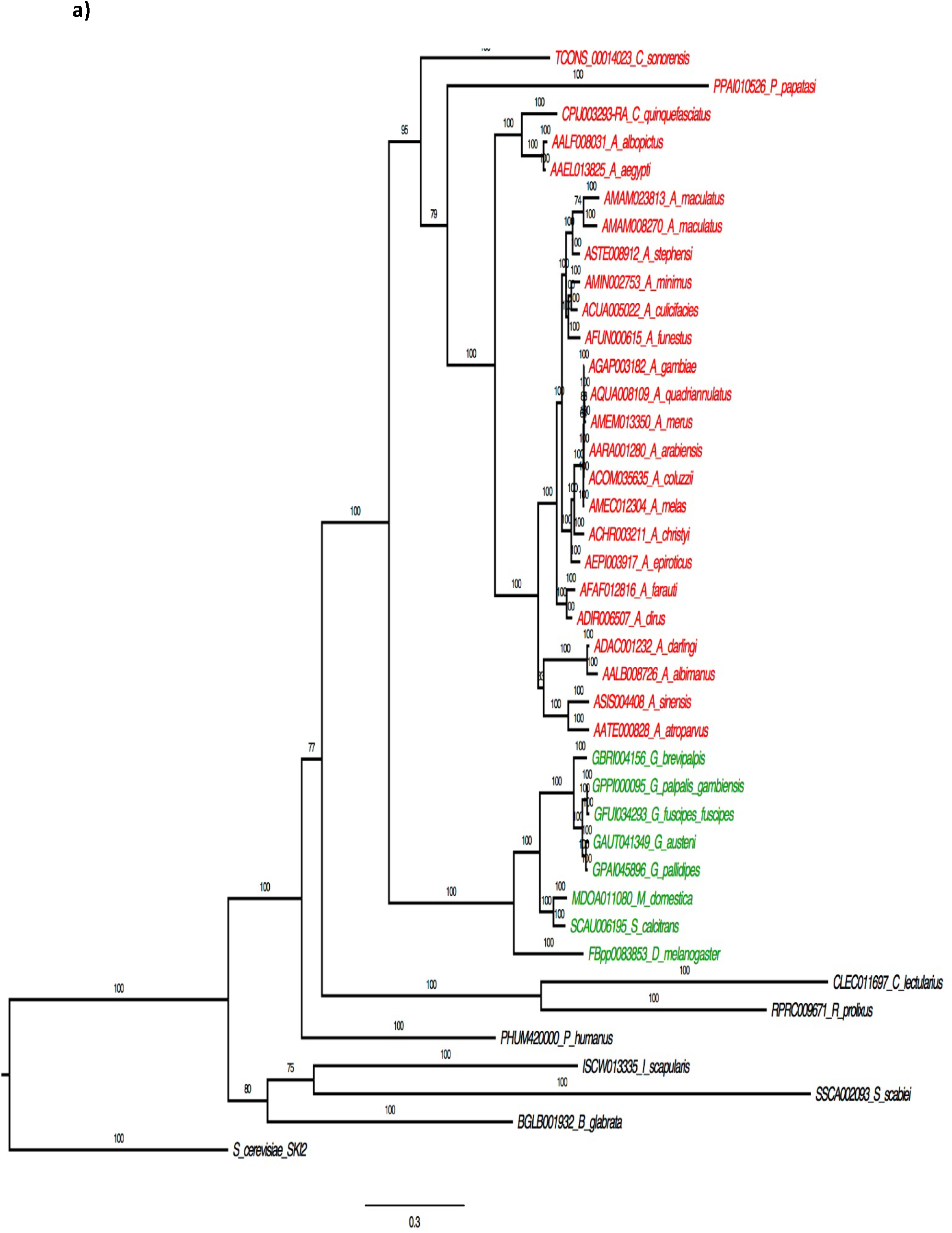
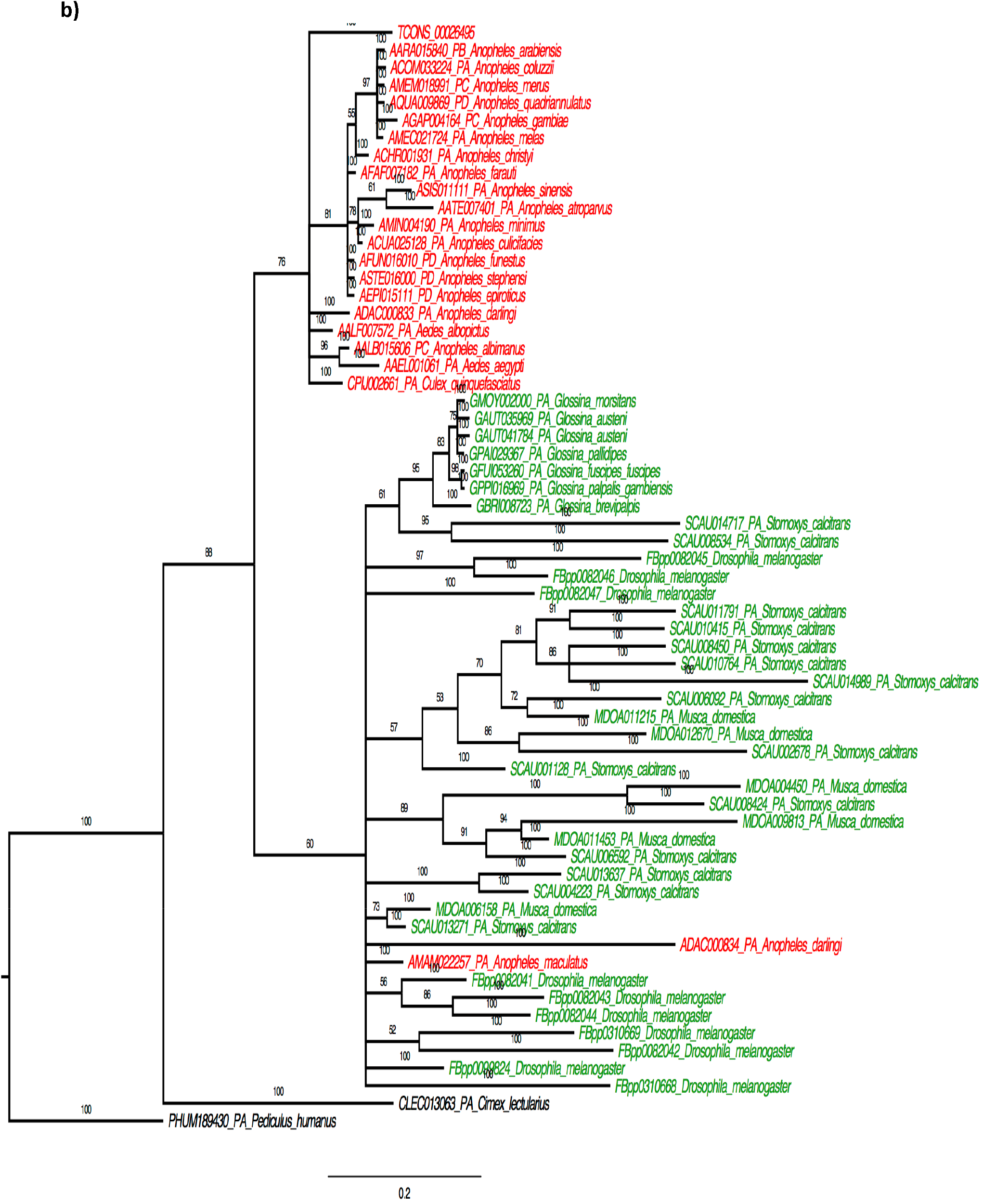
Bayesian phytogenies of *ski2* (a) and *gst-1* (b) genes. Tip labels correspond to the Ensembl Identifier followed by the species name (in the case of *C. sonorensis*, the number that precedes correspond to the RNA transcript); colouring corresponds to the Nematocera in red and Brachycera in green. Labels on branches show the posterior probability values.

In the *gst-1* gene from *C. sonorensis*, the most similar BLAST hit is that of *C. variipennis* (synonym of *C. sonorensis*; AAB94639.1) [25]. The second closest was the *gst-1* copy of *An. gambiae* (AGAP004164), for which 61 orthologues have been identified in 31 other species in VectorBase (Supplementary Table S6b). The mean evolutionary distance between the gst-1 protein of the Diptera species, estimated with the same model parameters as for ski2 protein, was 0.647. The mean distance between the gst-1 protein of *C. sonorensis* and that of mosquitoes was 0.199, while its evolutionary mean distance to Brachycera was 0.866. The pairwise distances ranged between 0.000 (observed between several sequences) and 2.914 (between the copy of *M. domestica* MDOA012670 and *An. coluzzi* ACOM0333224). The mean evolutionary distance within the Nematocera was 0.141, and 0.722 within the Brachycera. The phytogeny of the *gst-1* protein recovers the two main clades in Diptera, Brachycera and Nematocera, but there is little resolution within these clades (Fig. 3b).

### Immune response related genes in *C. sonorensis*

All *D. melanogaster* genes from FlyBase that had InterPro hits containing Toll, Imd and JAK/STAT associated to their names and all GO terms matching to the Toll, Imd and JAK/STAT pathway terms were used as reference to identify the homologues in *C. sonorensis* (Supplementary Table S7, and S8; Supplementary Figure S3). 36 of 42 genes identified in the Toll pathway of *D. melanogaster* were also identified in *C. sonorensis*; eight of the 11 genes of the Imd signalling pathway in *D. melanogaster* were identified in the *C. sonorensis* genome and 10 of the 13 *D. melanogaster* JAK/STAT pathway genes were identified. There are four instances in which the same *C. sonorensis* genes were identified as different immune-related genes, and we have verified them. Two *C. sonorensis* genes, CSONŬ12766 and CSONŬ15181, were predicted by the Compara workflow to be at the same time *Dorsal* and *Dif*. These two genes had 99.6% similarity at protein sequence level and 96% similarity at DNA sequence level, and are located in different scaffolds. The similarity to *D. melanogaster Dorsal* was 65% and to *Dif* was 61%. Four *C. sonorensis* genes, CSONOŬ1282, CSON001790, CSONOŬ7335, CSONOH712, were predicted to be both *Toll* (FBgn0262473) and *Tehao* (FBgn0026760) at the same time. Blastp searches identified all four genes as having leucine-rich repeats and Toll/Interleukin-1 receptor homology (TIR) domains, which suggests they could be identified as *Toll* rather than *Tehao*. However, with the similarity with *Toll* is about 3840% at the protein sequence level. Two *C. sonorensis* genes, CSON003218, CSON008584, have been identified as both *Peptidoglycan recognition protein LC* (Fbgn0035976) and *Peptidoglycan recognition protein LF* (Fbgn0035977). Using Blastp searches, the two genes showed partial similarity higher than 50% to several PGRPs at the protein level, not being able to differentiate between them using Blast. Finally, one single *C. sonorensis* gene (CSON004570) was identified as Unpaired 1, 2 and 3 (FBgn0004956, FBgn0030904, FBgn0053542). Using Blastp, the *C. sonorensis* protein had partial identities of 25-30% to these three Unpaired proteins ofíλ *melanogaster*. Similar to the cases above, it is not possible to identify the exact identity of the gene without further analyses.

Additional searches for immune related genes in *C. sonorensis* were carried out using BLASTp with Λe. *aegypti* and *Cx. quinquefasciatus* gene sequences from the ImmunoDB database (http://cegg.unige.ch/Insecta/immunodb) as queries (Supplementary Table S9 and S10). Four annotated genes in *C. sonorensis* were identified as potential antimicrobial peptides (AMPs) belonging to the gene subfamilies Attacin, Cecropin, Defensin and Diptericin (Supplementary Table S9a); however, the analyses did not identify more than a single gene within each of the AMP subfamilies. Eight potential Toll-like receptors were identified using BLASTp although there were three instances in which the most significant identities were different depending on the species used to provide input data (Supplementary Table S10), reflecting the large difference in sequence between the homologous genes in *Ae. aegypti* and *Cx. quinquefasciatus*. All five genes belonging to the Toll pathway (Supplementary Table S10) and all genes of the JAK/STAT signal transduction pathway were identified in *C. sonorensis* (Supplementary Table S10); while of the Imd pathway, one gene was identified in five of the eight subfamilies (Supplementary Table S10).

The Compara approach reconciles the gene tree (derived from sequence similarity metrics) with the species tree to identify likely orthologous relationships; but requires an initial clustering step (which can lead to the inclusion of sequences into the wrong trees) [45]. The only instance in which the two approaches identify a different *C. sonorensis* gene and the orthologue of a gene in the above pathways is that of *Pelle* (which activates the Toll Receptor), for which BLAST identifies CSON012655 as its homologue, and Compara identifies CSON013584 and CSON013585. There were other cases in which BLAST identifies a potential homologue while Compara does not (e.g. TNF-receptor-associated factor 6, Trafő, or tube) and genes Imd and Fadd, neither of the two approaches identify a homologue in *C. sonorensis*.

It should be noted that the results presented in this study come from automated pipelines. Although some cases have been manually verified, there are still genes which have not been identified in this study and other that need improvement in their annotation. It is expected that the annotation and curation of the genome will advance as it is used by the research community.

## Discussion

This study has produced the first *de novo* genome assembly of the Dipteran family Ceratopogonidae, which includes important vectors of emerging and re-emerging veterinary arboviruses [8, 65]. The closest relative to this group for which a genome of comparable quality is available is the non-biting midge, *Polypedilum vanderplanki* Hinton (Family: Chironomidae) [66], which diverged from the Ceratopogonidae some 220 million years ago. This study provides a primary resource for comparative studies with other arbovirus vector species (e.g. mosquitoes [67]). It also will facilitate efforts to produce transgenic *Culicoides* to control *Culicoides*-borne pathogens, although the limitations on colony production for the other epidemiologically relevant species of *Culicoides* remains a major constraining factor [14, 68].

The genome assembly of *C. sonorensis* is not wholly contiguous, however, its general features are consistent with similarly sized genomes from other species. Thus, features such as the repeat content, gene count and level of alternative splicing are within the ranges observed in *Ae. aegypti, An. gambiae* and *D. melanogaster*. Coverage of well-conserved gene families is complete and comparable with other published assemblies (Supplementary Table S12). Various measures (the high proportion of BUSCO genes present in multiple copy, and estimates of genome size and redundancy using read mapping and alignment approaches) indicate the presence of redundancy in the assembly (Supplementary Table S13), which could result from genetic variation within and amongst individuals from which sequenced material was extracted. However, the individuals come from a colony that has not had any outbreeding for over 40 years, which should have reduced the genetic variation among the individuals of the colony. Long read-sequencing approaches would likely help reduce this redundancy. However, the gene set has been annotated with a comparable number of InterPro domains and high-level GO terms as *An. gambiae* and other Diptera, while the number of specific GO terms is comparable to other genomes that have not been extensively manually annotated with functional descriptions. The molecular functions of ion binding, oxidoreductase activity, kinase activity, hydrolase activity acting on glycosyl bonds, methyltransferase activity all highly represented, as is the tRNA metabolic process. In addition, the genes of the immune-related pathways have been identified and annotated, revealing complete or nearly-complete orthologous pathways. Overall, these observations indicate that the present assembly and annotation of the *C. sonorensis* genome represents a reliable resource to use in genetic studies of biting midges. Nevertheless, it is still highly fragmented and the use of long-read sequencing technologies like PacBio would help improving the scaffolding. In addition, the use of this genomic resource by the community will help improve the annotation of the genes. This is the case for many of the genomes of vector species that have been neglected over the years.

A striking feature of the *C. sonorensis* genome is its GC content, which is among the lowest reported for Dipteran species (although close to identical to the non-biting midge *P. vanderplanki* [66] and other non-Dipteran arthropods like the pea aphid, *Acyrthosiphon pisum* Harris [69]). This is not just a feature of the repeated content, but extends into the protein-coding genes (which have a higher GC content than the noncoding regions, but less than in comparable species: Table 1). Recent comparative analyses of codon usage in Diptera and Hymenoptera showed a significant association between codon bias and high GC content in Diptera, but low GC content in Hymenoptera [70]. Thus, the low GC content observed in *C. sonorensis* could reflect a codon bias in comparison to other Diptera species. Functional investigations of GC content have also revealed that it has an impact on genome functioning and species ecology in microbes [71], vertebrates [72] and plants [73]. It is also known that the GC content has an impact in the efficiency of sequencing technologies [74], Regions with low GC content have less coverage than more GC balanced ones. This characteristic of the *C. sonorensis* genome should be accounted for in future genomic studies, and appropriate experimental designs that take into account the low GC content should be used [e.g. 75].

Studies investigating gene expression during infection with BTV identified four candidate genes whose regulation was correlated with vector competence, despite only using a single time-point at eight days post-infection and pools of *C. sonorensis* exhibiting different degrees of virus dissemination. Interestingly, the elevated expression of *gst-1* in refractory *Culicoides* observed in this study is consistent with a previously proposed genetic mechanism for *C. sonorensis* vector competence for BTV [13, 15, 24], These studies identified a locus controlling vector competence in the AA colony, the origin colony of PIR-s-3 colony used in the present study. The former study used a maternally inherited 90kd protein consistent with genetic studies that suggested maternal inheritance of the competence controlling factor identified in unfertilized eggs of refractory females to isolate a clone that upon sequencing was identified as *gst-1* [25]. Though the current study evaluated gene expression after exposure to BTV and the previous study compared genetically selected resistant and susceptible families using individuals from these families who were not exposed to BTV, both independently identified involvement of *gst-1*. Thus, the differential expression of the gene in the current study is confirmation of its involvement in BTV infection in AA colony, although its role requires functional validation.

From the evolutionary perspective, the phytogenies of ski2 and gst-1 reflect the accepted evolutionary history of the included Diptera species, although the phylogenetic tree of ski2 shows more resolution within Nematocera and Brachycera than the gst-1. It is also interesting to note the two contrasting modes of evolution of gst-1 between the two main clades of Diptera. The gst-1 protein in the Brachycera clade shows multiple duplications in comparison with the Nematocera and the estimated diversity in the Brachycera is higher (0.722) than that in the Nematocera (0.141). Within the Nematocera the level of diversity of the gst-1 protein is tower than in ski2.

Future studies of vector competence in *C. sonorensis* must characterize genes that facilitate BTV competence and differentiate these from differentially expressed genes that are the consequence of BTV infection. To determine variation in the mechanism and factors controlling BTV-vector competence in the AA and PIR-s-3 colonies observed in the present study, it will be necessary to carry out studies of different populations of *C. sonorensis* and other *Culicoides* vectors and non-vector species. Once controlling genes are identified it will be essential to assess the effect of polymorphisms, the various functions of the identified genes in the absence of the pathogen to evaluate factors that influence their frequencies, and compare these processes between different species of *Culicoides* [23].

The present paper presents the genome of *C. sonorensis* and its use in transcriptomic analyses to provide information that will help elucidate the transmission of viruses by vectors and provide new avenues of research to understand vector competence for BTV. We present a list of candidate genes that will be further explored to show their involvement in the transmission of BTV by *C. sonorensis*. Furthermore, the results also provide evidence for the involvement of genes belonging similar functional families in response to virus infection in different species belonging to different families of Diptera. These results show the importance of comparative analyses to interrogate the evolution of host-pathogen interactions, and the *C. sonorensis* genome will facilitate such studies.

## Conclusions

Here we present the first annotated genome of the biting midge, *C. sonorensis*, a vector of economically important viruses of livestock, such as Bluetongue virus (BTV) and Schmallenberg virus. This genome has been fundamental in gaining information about the genetics of BTV transmission. Thus, gene expression comparison between females susceptible to a full infection by BTV and those females refractory to full BTV infection, identified 165 genes that are candidates to be involved in vector competence in this species. Of these genes, the *gst-1* gene was identified previously to be involved in BTV transmission, while the antiviral helicase *ski2* involved in suppressing the replication of dsRNA viruses. Overall, the publication of the genome and transcriptomes will lead to advances in the control of economically relevant arboviruses of livestock. In addition, this genome fills a phylogenetic gap of more than 200 million years, and will be a useful source of comparative data for studies of vector competence across Diptera.

## Declarations

### Ethics approval and consent to participate

Not applicable.

### Consent for publication

Not applicable.

### Availability of data and material

All sequencing data, genomic and transcriptomic, generated in this study has been deposited at the European Nucleotide Archive (ENA) database under the accession number PRJEB19938.

### Author’s contributions

MF, SC and PK conceived the research project RMH, MH, IMA, RS, LEH, AGU, EV and LC have performed the laboratory experiments and bioinformatic analyses. DN and CS have helped with the annotation of the immune-related genes. RMH, MF and SC have written the manuscript and PK, WT, ND and CS have provided insightful revision. All authors read and approved the final manuscript.

## Acknowledgements

We thank Eric Denison and James Barber in the insectary for their supply of reagents, cell lines and insects for this study. We would like to thank the three reviewers and the editor for their positive comments and suggestions.

## Supporting Information Legends

### Supplementary Material Tables

**Table S1.** Differentially expressed genes between vector competent and vector refractory *C. sonorensis* females identified using CummeRbund. The table shows the fpkm values for each phenotype, the log2 fold change of expression levels and the significance level of the change. Genes that are not annotated in the *C. sonorensis* genome are denoted with the name XLOC_XXXXXX. Genes belonging to an enriched GO term are shown in bold.

**Table S2.** BLAST results for the differentially expressed genes between vector competent and refractory transcriptomes showing the top hit as determined by BLAST.

**Table S3.** Ranking of the expression stability of the reference genes tested for use in the quantitative reverse transcription PCR (RT-qPCR). Three different tests were used to rank the expression stability of the genes, BestKeeper, gNorm and NormFinder. *r*-correlation coefficient, Cq – quantification cycle, SE – standard error, SD – standard deviation.

**Table S4.** Primers forward (F), reverse (R) and hydrolysis probe (P) sequences used for RT-qPCR analysis of the expression levels of *ski2, gst, gst-1* and *Toll*-like.

**Table S5.** Difference of expression levels in four genes (*Toll*-like, *gst, gst-1* and *skiZ*) between vector-competent and vector-refractory females assessed by RNAseq and RT-qPCR. * denote a difference in expression between refractory and competent females with a *P* < 0.05 (RT-qPCR significance estimated using a Welch Two Sample t-test with d.f. = 1).

**Table S6.** Orthology analysis of two of genes *Ski2* **(a)** and *gst-1* **(b)**.

**Table S7.** Immune pathways genes identified in *C. sonorensis* using the Ensembl Compara pipeline and the *D. melanogaster* genes as reference, **a)** Toll pathway; **b)** Imd pathway; **c)** Jak/Stat pathway.

**Table S8.** Homologue immune genes in several invertebrate species identified using *D. melanogaster* genes as reference all with InterPro and GO terms associated with Toll **(a)**, Imd **(b)** and Jak/Stat **(c)**, in the Ensembl Compara pipeline.

**Table S9.** Immune related genes identified in *C. sonorensis* using BLASTp top hits with the gene copies from *Ae. aegypti* and *Cx. quinquefasciatus* available in the ImmunoDB database (http://cegg.unige.ch/Insecta/immunodb) as query, a) Anti-Microbial Peptides; b) Toll receptors; c) Toll path; d) jak/Stat path; e). Imd path. * - Top BLASTp hits different found whence, *aegypti* and *Cx. quinquefasciatus* were used as query.

**Table S10.** Details of the BLASTp results using/le. *aegypti* (a) and *Cx. quinquefasciatus* (b) as query to identify the *C. sonorensis* genes of the immune pathways.

**Table S11.** Read mapping and estimate of assembly size, by read mapping using BWA, according to the method of Schell et al. [36].

**Table S12.** BUSCO analysis using the Insecta data set (version 3.0.2) against the assembled genome sequence of six species, including *Culicoides sonorensis*.

**Table S13.** Comparison of the original, and redundancy-reduced, assemblies of C. *sonorensis*.

**Table S14.** Number of distinct InterPro and GO terms annotated to annotated proteins of *C. sonorensis* and six other Diptera species.

**Table S15.** Tandemly repeated gene arrays in *C. sonorensis* versus other insect species. Longest gene arrays in *C. sonorensis* with matching InterPro domain architecture and maximum 100,000 nucleotide between successive members. The longest array with the same domain signature from 3 other insect species is given for comparison.

### Supplementary Material Figures

**Figure S1.** Distribution of C_q_ values from RT-qPCR used to detect BTV-1 virus infection in the two virus-feeding experiments (in blue and orange) of *C. sonorensis*. The vertical, dashed line corresponds to a C_q_ value of 27 used to separate vector competent from refractory females.

**Figure S2.** Classification and functional distribution of the genes differentially expressed between vector competent and refractory *C. sonorensis* per the Gene Ontology level 6. Blue: Molecular Function; Green: Cellular Component; Pink: Biological Process.

**Figure S3.** Number of genes in the Toll (a), Imd (b) and Jak/Stat (c) pathways that have been duplicated or lost in different species of Diptera as identified using the Ensembl Compara pipeline.

### Supplementary Material Methods

#### DNA extraction

Protocol followed to extract DNA from *C. sonorensis* samples.

#### Genome annotation and comparative analysis

Details about the genomes of species used in the gene annotation and in the comparative analyses.

#### RNA extraction

Protocol used to extract RNA from *C. sonorensis*.

#### RT-qPCR for the detection of BTV-1 infection

Description of the method used to detect the presence of BTV-1 in *C. sonorensis* females after the experimental infections.

#### RT-qPCR protocol for the validation of differentially expressed genes

Description of the protocol used to verify the change in expression of *Toll-like protein, glutathione-s-transferase, glutathione-s-transferase 1* and *ski2 antiviral helicase* genes observed in the transcriptomes of vector-competent and refractory samples.

#### Alignment of gst-1 and ski2

The commands used to align the protein sequences of gst-1 and ski2. These alignments were then used to run the phylogenetic analyses.

